# Multivariate analysis of the cotton seed ionome reveals a shared genetic architecture

**DOI:** 10.1101/213777

**Authors:** Duke Pauli, Greg Ziegler, Min Ren, Matthew A. Jenks, Douglas J. Hunsaker, Min Zhang, Ivan Baxter, Michael A. Gore

**Author notes:** Corresponding author: Michael A. Gore, 358 Plant Science Building, Cornell University, Ithaca, NY 14853. Present address: School of Plant Sciences, University of Arizona, Tucson, AZ 85721, USA. **Author contributions:** D.P. and M.A.G. co-wrote the manuscript; D.P. led data analysis; G.Z. and I.B. conducted elemental profiling and consulted on data analysis; M.R. and M.Z. carried out genetic mapping; M.A.J. oversaw soil and nutrient management; D.J.H. managed irrigation scheduling; M.A.G. overall project management, design, and coordination, oversaw data analysis; all authors contributed to the critical review of the manuscript. **Funding information** This research was supported by Cotton Incorporated Fellowship (D.P.), Cotton Incorporated project funds 13-580 (I.R.B.) and 14-194 (M.A.G), Cornell University startup funds (M.A.G), United States Department of Agriculture - Agricultural Research Service (USDA-ARS) (D.H., I.B. and G.Z.), and National Science Foundation IOS-1238187 (M.A.G).

## Abstract

To mitigate the effects of heat and drought stress, a better understanding of the genetic control of physiological responses to these environmental conditions is needed. To this end, we evaluated an upland cotton (*Gossypium hirsutum* L.) mapping population under water-limited and well-watered conditions in a hot, arid environment. The elemental concentrations (ionome) of seed samples from the population were profiled in addition to those of soil samples taken from throughout the field site to better model environmental variation. The elements profiled in seeds exhibited moderate to high heritabilities, as well as strong phenotypic and genotypic correlations between elements that were not altered by the imposed irrigation regimes. Quantitative trait loci (QTL) mapping results from a Bayesian classification method identified multiple genomic regions where QTL for individual elements colocalized, suggesting that genetic control of the ionome is highly interrelated. To more fully explore this genetic architecture, multivariate QTL mapping was implemented among groups of biochemically related elements. This analysis revealed both additional and pleiotropic QTL responsible for coordinated control of phenotypic variation for elemental accumulation. Machine learning algorithms that utilized only ionomic data predicted the irrigation regime under which genotypes were evaluated with very high accuracy. Taken together, these results demonstrate the extent to which the seed ionome is genetically interrelated and predictive of plant physiological responses to adverse environmental conditions.

**One sentence summary:** The cotton seed ionome has a shared genetic basis that provides insight into the physiological status of the plant.

## INTRODUCTION

Plant growth, development, and survival are highly dynamic processes that can be altered by a myriad of favorable and adverse environmental conditions including abiotic stresses such as water deficit and high temperature. The physiological responses of plants to changing environmental conditions are difficult to observe and quantify at the population level, thus unraveling the genetic mechanisms responsible for the variability of these traits remains a formidable challenge. This is especially true for understanding plant nutrient and mineral uptake, which occurs below ground and is obscured by the soil environment. Because of the challenges associated with phenotyping below-ground traits such as root architecture and resource capture, there exists a gap in our knowledge about these fundamental biological mechanisms.

Elemental uptake is a critical function driven by physiological and biochemical processes occurring throughout the life cycle of a plant. Many factors affect elemental accumulation including availability in the soil environment, bioavailability within the plant, and the ability of the plant to mobilize and translocate them throughout cells and tissues. Additionally, physiological parameters such as root depth, permeability of root barriers, and rate of transpiration all affect the capacity of elements to enter and move throughout the plant. As a result, the elemental content and composition (the ionome) of plant tissues such as leaf and seed can be considered a “readout” of the summation of these processes and thus provide insight into plant stress response (Salt et al., 2008).

Plant ionomic analysis was originally developed in Arabidopsis, where it was used to identify elemental accumulation mutants and characterize their physiological responses to the environment (Lahner et al., 2003). This approach, wherein the elemental composition of plant tissues is described and useful genetic mutants identified, has been extended to a variety of species including rice (Norton et al., 2010; Zhang et al., 2014; Pinson et al., 2015), maize (Baxter et al., 2013; Baxter et al., 2014; Mascher et al., 2014; Gu et al., 2015; Asaro et al., 2016), barley (Wu et al., 2013), soybean (Ziegler et al., 2013; Huber et al., 2016), tomato (Sánchez-Rodríguez et al., 2010), and other crops (Chen et al., 2009; White et al., 2012; Shakoor et al., 2016). These studies have shown that elemental traits are heritable and thus amenable to genetic mapping, but they have also demonstrated that individual elements exhibit phenotypic correlations that could arise from shared genetic control, overlap in membrane transporters at the cellular level, common physiochemical properties in the soil and rhizosphere, macroscale environmental conditions, or some combination of these factors. One such set of factors that may have an impact on the ionome are heat and drought stresses, environmental conditions that not only affect the plant itself through water availability but also through modification of the soil environment from which nutrients are acquired (Vietz, 1972).

Heat and drought are two of the most common abiotic stresses that occur simultaneously in agricultural production areas, often with devastating effects on yield and economic returns (Rizhsky et al., 2002; Fannin, 2012). At the cellular level, these stresses affect photosynthesis through adverse regulation of stomata and resulting CO2 uptake, thereby decreasing whole plant health and fitness (Chaves et al., 2003; Taiz and Zeiger, 2006; Prasad et al., 2008). Increased variability in weather patterns could have a significant impact on crop yields, as well as threaten the manufacture of critical bio-based commodities in agricultural areas already at risk (Wheeler and von Braun, 2013; Thornton et al., 2014). Expanding the understanding of how plants respond to abiotic stress at the physiological level has the potential to help optimize breeding strategies focused on improving crop stress resilience. Nowhere is this truer than for cotton (*Gossypium* spp.), a crop with no naturally occurring substitute that can be produced on the scale demanded by economic markets.

Cotton is the most widely grown fiber crop in the world, being produced in over 80 countries and responsible for a multi-billion dollar industry. In 2016, 21 million metric tons of cotton fiber were produced globally (Cotton Inc, 2017). In the US, the largest global exporter of cotton, the annual value of the crop is over $5 billion, translating into $25 billion generated in value-added products and services (USDA Economic Research Service, 2015). Currently, 65% of US cotton acreage is produced under rainfed agricultural systems, with global data reflecting similar conditions in other countries (National Cotton Council of America, 2015). Because of this, cotton, like all crops, is threatened by the effects of climate change including decreased rainfall, increased temperatures, and highly variable weather patterns (Dabbert and Gore, 2014). To contend with temperature and precipitation changes that are unfavorable to crop growth, new technologies such as field-based, high-throughput phenotyping have been investigated to support the more efficient and effective development of stress-resilient cultivars (Thorp et al., 2015; Pauli et al., 2016a; Pauli et al., 2016b). Ionomic profiling, which can be done on seed, is a complementary technology that could provide insight into the physiological status of the plant in relation to its growing environment.

To make progress towards this goal, we assayed the elemental profiles of seed in an upland cotton (*G. hirsutum* L.) recombinant inbred line (RIL) mapping population that was evaluated under two contrasting irrigation regimes over three years. The 14 elements that were profiled were arsenic (As), calcium (Ca), cobalt (Co), copper (Cu), iron (Fe), potassium (K), magnesium (Mg), manganese (Mn), molybdenum (Mo), nickel (Ni), phosphorus (P), rubidium (Rb), sulfur (S), and zinc (Zn). Ionomic analyses were also conducted on soil samples collected from multiple depths throughout the field site where the population was grown in order to investigate relationships between soil and seed elemental concentrations. Complementary quantitative trait loci (QTL) mapping methods, consisting of Bayesian classification and frequentist multivariate approaches, were employed to identify regions of the cotton genome controlling phenotypic variation for elemental concentrations. To test whether the ionome could serve as an indicator for environmental growing conditions, various supervised machine learning methods were implemented to predict the irrigation regime under which RILs were grown using only the ionomic data. Empirical results of this study demonstrate that the ionome is a dynamic system that responds in a coordinated manner due to its shared genetic architecture, providing valuable information on the physiological status of plants.

## MATERIALS AND METHODS

### Plant Material and Experimental Design

The plant material and experimental design have been extensively described in Pauli et al. (2016a). Briefly, 95 recombinant inbred lines (RILs) from the TM-1×NM24016 biparental mapping population (Percy et al., 2006; Gore et al., 2012) and commercial check cultivars were evaluated at the Maricopa Agricultural Center (MAC) of the University of Arizona located in Maricopa, AZ, in 2010-12. The 95 RILs, parental lines, and commercial cultivars were grown under two irrigation regimes, water-limited (WL) and well-watered (WW), at a field site of predominantly sandy clay loam soil texture. Each year, the trial was arranged in an alpha (0, 1) lattice design with two replications per irrigation regime. Plots were one-row measuring 8.8 m in length with 1.02 m inter-row spacing, and thinned to a density of ~4.1 plants m^−2^. The trial was managed with conventional cotton cultivation practices. Meteorological data were recorded by an automated Arizona Meteorological Network (AZMET) weather station (ag.arizona.edu/azmet/index.html) located 270 m from the field site (Brown, 1989).

To establish the crop, several furrow irrigations were applied during the first 10-14 days after planting, then subsurface drip irrigation (SDI) was used for the remainder of the field season. In late May of each year after plant establishment, neutron moisture probe access tubes were installed to a depth of 1.6 m at 56 selected locations throughout the field with an equal number of tubes in the WL and WW treatment plots. Weekly soil water content measurements in 0.2 m increments from a depth of 0.1 to 1.5 m were made for all probe locations from early June through early October in each year using field-calibrated neutron moisture probes (Model 503, Campbell Pacific Nuclear, CPN, Martinez, CA). The scheduling of the WW SDI treatment was performed using a daily soil-water-balance model calculated over the cotton root zone as previously described in Andrade-Sanchez et al. (2014). Soil-water-balance model inputs included estimated daily crop evapotranspiration (ETc) as determined from FAO-56 dual crop coefficient procedures (Allen et al., 1998), metered irrigation depths, and precipitation data from the AZMET weather station. For ETc, the cotton basal crop coefficient (Kcb) values were 0.15, 1.2, and 0.52 for the initial, mid-season, and end of season values, respectively. These parameters were constructed into an FAO-56 Kcb curve using the growth stage lengths developed locally by Hunsaker et al. (2005) for a typical 155 day cotton season. Additional crop and soil parameters used in calculating the daily soil water balance were as those presented in Hunsaker et al. (2005; table 3), with the exception of the fraction of soil wetted by irrigation, which was reduced to 0.2 for the SDI.

Irrigations to the WW plots were applied to refill the root zone water content to field capacity when approximately 35% of the available soil water had been depleted. Starting mid-July, the WL plots received one half of the irrigation amounts applied to the WW plots. The WL irrigation regime was imposed when more than 50% of the plots were at first flower to minimize the interaction of phenology and soil moisture deficit. The weekly soil water content measurements for the WW treatment were used to monitor the actual soil water depletion and adjust the modeled soil-water-balance depletion when needed.

### Soil and Seed Sampling for Ionomic Profiling

In 2010 and 2012, soil samples were collected from the 56 neutron access tube sites at five depth intervals: 0 – 30; 30 – 60; 60 – 90; 90 – 120; and 120 – 150 cm. The locations of neutron tubes for 2011 were the same as for 2010, thus soil sampling was not repeated in 2011. In 2012, the neutron probes were redistributed in the field and soil samples were taken from the new neutron access tube sites. Soil sampling was performed within each of the five depth intervals at a probe location (five samples per 56 probe locations). The collected samples were homogenized and then sent to the Donald Danforth Plant Science Center (DDPSC) where they were dried and then ground to obtain a soil particle size less than 4 mm. From these prepared soil samples, two 50 mg subsamples were taken per each depth interval within a probe location and placed in digestion tubes for ionomic profiling.

Prior to mechanical harvest at the end of the season, 25 bolls were harvested from each experimental plot and processed using a laboratory 10-saw gin. From the processed boll samples, six seeds were randomly sampled and sent to DDPSC where they were individually weighed and placed in digestion tubes for further processing.

### Determination of Elemental Composition by ICP-MS

Samples for both soil and unground seed were digested, analyzed by inductively coupled plasma mass spectrometry (ICP-MS), and data corrected for loss of analyte during sample preparation and drift as described in Ziegler et al. (2013). Briefly, both soil and unground seed were separately digested in 2.5 mL nitric acid containing 20 parts per billion (ppb) indium as a sample preparation internal standard at room temperature overnight, then heated to 100 °C for 3 h and diluted to 10 mL using ultra-pure water (UPW). The nitric acid digestion of soil samples is a partial digestion procedure. However, nitric acid digestion is sufficient to provide an adequate sample of elemental content available for biological uptake. Samples were diluted in-line with 5x volume of UPW containing yttrium as an instrument internal standard using an ESI prepFAST autodilution system. Elemental concentrations were measured using a Perkin Elmer Elan 6000 DRC-e mass spectrometer for all seed samples and the 2012 soil samples. A Perkin Elmer NexION 350D with helium mode enabled for improved removal of spectral interferences was used to measure elemental concentrations of the 2010 soil samples. Instrument reported concentrations were corrected for the yttrium and indium internal standards and a matrix matched control (either pooled soil or pooled seed sample). The control was run every 10 samples to correct for element-specific instrument drift. The same control was used in every ICP-MS run to correct for run-to-run variation.

To correct seed elemental concentrations for weight and run-to-run variation, we followed the method of Asaro et al. (2016). Briefly, elemental concentration was regressed against sample weight and ICP-MS run. The residuals from this model reflected the deviance of samples from the population mean and were used as the weight-normalized phenotype. Because a uniform amount of soil was digested for each sample, this normalization technique was not necessary for the soil samples. Therefore, soil concentrations were simply converted to parts per million, ppm, (mg analyte/kg sample) by dividing instrument reported concentrations by the 50 mg sample weight.

Analytical outliers for seed samples were removed first using the non-weight-normalized values and again after normalization. Outliers were identified by analyzing the variance of the six seed replicate measurements and excluding an elemental measurement from further analysis if the median absolute deviation (MAD) was greater than 6.2. After outlier removal and weight-normalization, the elemental concentrations were transformed to non-negative by adding a constant to every sample so that the smallest value for each element quantified had a value greater than 10. This was done to avoid boundary constraint issues during variance component estimation. Final concentrations were calculated by taking the mean of the six individual seed elemental concentrations and reported as parts per billion, ppb.

### Soil Ionomic Data Analysis and Spatial Interpolation

To identify and remove significant outliers from the raw soil ionomic data, we fitted a mixed linear model for each element in ASReml-R version 3.0 (Gilmour et al., 2009). For the elements calcium (Ca), arsenic (As), and iron (Fe), soil sample data were log transformed to stabilize variances based on preliminary analyses using mixed linear models. The full model (Equation 1) fitted to the data was as follows:

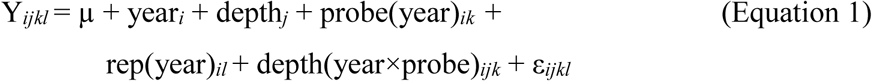

in which Y_*ijkl*_ is an individual soil sample observation; μ is the grand mean; year_*i*_ is the effect of the *i*th year; depth_*j*_ is the effect of the *j*th soil sample depth; probe(year)*ik* is the effect of the *kth* neutron probe site nested within the *i*th year; rep(year)_*il*_ is the effect of the *l*th technical replication of the homogenized soil sample taken from a neutron probe site nested within the *i*th year; depth(year×probe)*ijk* is the effect of the *j*th soil sample depth level nested within the *kth* neutron probe site and *i*th year; and ε _*ijkl*_ is the random error term following a normal distribution with mean zero and variance *σ*^2^. The model terms year *i* and depth *j* were fitted as fixed effects while all others were fitted as random effects. To detect significant outliers, Studentized deleted residuals (Neter et al., 1996) were used with degrees of freedom calculated using the Kenward-Rogers approximation (Kenward and Roger, 1997).

Once all outliers were removed for each element, an iterative mixed linear model fitting procedure of Equation 1 was conducted in ASReml-R version 3.0 (Gilmour et al., 2009). Likelihood ratio tests were conducted to remove all terms fitted as random effects from the model that were not significant at α = 0.05 (Littell et al., 2006) to generate a final, best fitted model for each element. This final model was used to generate best linear unbiased predictors (BLUPs) for each unique neutron probe location for 2010 and 2012. Sequential tests of fixed effects were carried out with degrees of freedom calculated using the Kenward-Rogers approximation. For those elements in which log transformation was required (As, Ca, and Fe), BLUPs were back transformed prior to further analyses.

The calculated BLUPs for each neutron probe location within the field were then utilized for spatial interpolation using geostatistical methods. Conventionally in geostatistical analyses, an estimate of the variogram model is calculated from the observed data points that describe the spatial variability of the underlying processes in the given physical area of study. The derived model accounting for the spatial relationship between sampling locations, which describes the covariance as a function of distance between points (Yates, 1948), is then used to predict the values at unsampled but known locations – this methodology is known as kriging (Webster and Oliver, 2007). Due to the subjective nature of deriving empirical variograms, we used an iterative model fitting procedure to obtain initial model parameters using the *automap* package (Hiemstra et al., 2009) in R (R Core Team, 2016). The BLUPs for elements As, Ca, Fe, K, Mg, and P were log transformed to stabilize variances for model fitting. Through implementation of the ‘autoKrige’ function in *automap*, eight spatial relationship models (spherical, exponential, Gaussian, Matern, Bessel, circular, pentaspherical, and Stein’s parameterization of Matern) were tested, and for each spatial model used, seven values of the range parameter (the distance at which spatial dependency is no longer present, values of 10, 15, 20, 25, 30, 35, and 40 m) were iterated over to find the optimal value. All other parameters (e.g., nugget variance and sill) were calculated by the software. To assess the model fit for each combination of spatial model and range distance, the sums of squares error was extracted from the fitted model to determine reasonable starting values for all model parameters.

In the next step, the complete set of estimated parameters (range, sill, nugget, bin width, and spatial model) from the best fitted model were used as starting values to generate an initial empirical variogram using the function “variogram” in the R package *gstat* (Pebesma, 2004). The generated empirical variogram and estimated parameters were then passed to the auto-fitting function “fit.variogram” for further model optimization and parameter estimation with sample point weightings determined by *Nj/h^2^j*, where *N* is the number of point pairs and *h* is the distance between points. The final model for each element was then assessed by visual inspection to ensure they were adequately capturing the spatial variance structure and that the model fitting procedure selected optimal parameters. The optimized models for each element were then used to conduct block kriging whereby element concentrations were predicted at the plot level for each georeferenced plot across the experimental field site for the three years.

### Seed Ionomic Data Analysis

The processing of the seed ionomic data was similar to that of the soil ionomic data. Initially, we fitted mixed linear model for each element in ASReml-R version 3.0 to identify and remove significant outliers from the raw data. Unlike the soil ionomic data, no transformation of elemental concentrations was required due to the weight normalization step described above. The full model (Equation 2) fitted to the data was as follows:

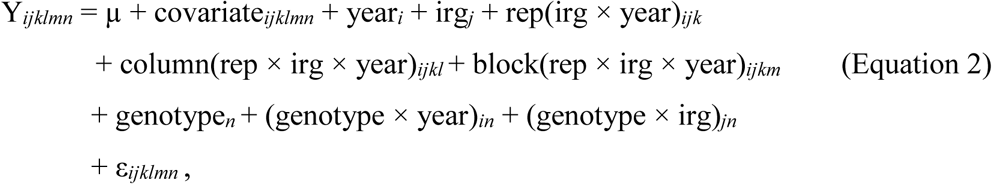

in which Y_*ijklmn*_ is the elemental concentration observation representing the adjusted model residual from the weight normalization step; covariate_*ijklmn*_ represents either flowering time (i.e., date of first flower, Julian calendar) or the interpolated soil element concentration for each observation; μ is the grand mean; year_*i*_ is the effect of the *i*th year; irg_*j*_ is the effect of the *j*th irrigation regime (WW or WL); rep(irg × year)_*ijk*_ is the effect of the *k*th replication within the *j*th irrigation regime within the *i*th year; column(rep × irg × year)_*ijkl*_ is the effect of the *l*th plot grid column within the *k*th replication within the *j*th irrigation regime within the *i*th year; block(rep × irg × year)*ijkm* is the effect of the *m*th incomplete block within the *k*th replication within the *j*th irrigation regime within the *i*th year; genotype*n* is the effect of the *n*th genotype; (genotype × year)*in* is the interaction effect between the *n*th genotype and the *i*th year; (genotype × irg)*jn* is the interaction effect between the *nth* genotype and the *j*th irrigation regime; and ε _*ijklmn*_ is the random error term following a normal distribution with mean zero and variance *σ*^2^. The two covariates, flowering time and soil element concentration, were tested individually for their significance at α = 0.05, and if found to be nonsignificant, they were removed from the model. The model terms genotype*n*, irg*j*, and (genotype × irg)*jn* were fitted as fixed effects, while all the other terms were fitted as random effects. To detect significant outliers, Studentized deleted residuals (Neter et al., 1996) were used with degrees of freedom calculated using the Kenward-Rogers approximation (Kenward and Roger, 1997).

Once outliers had been removed, iterative model fitting was conducted as described above with nonsignificant random terms removed from the model at a threshold of α = 0.05. The best fitted model was then used to generate both an overall (across three years) best linear unbiased estimator (BLUE) and a within year BLUE for each separate irrigation regime. Tests of model fixed effects were conducted as described for the soil element analysis.

For each element, broad-sense heritability on an entry-mean basis (*Ĥ^2^*) was estimated within each irrigation regime by reformulating Equation 2 to remove the irrigation regime term. Next all terms were fitted as random effects in order to estimate their respective variance components; however, if the covariate was found to be statistically significant it was retained in the model as a fixed effect. For each element, the variance component estimates from each final model were used to estimate *Ĥ^2^* (Holland et al., 2003) as follows:

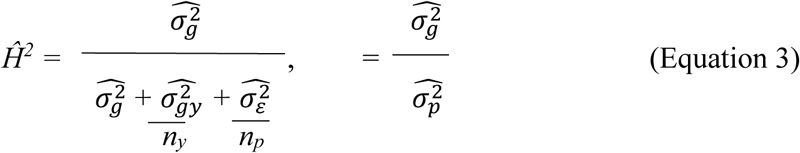

where 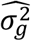 is the estimated genetic variance, 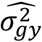 is the estimated variance associated with genotype-by-year variation, 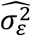 is the residual error variance, *n*_*y*_ is the harmonic mean of the number of years in which each genotype was observed, and *np* is the harmonic mean of the number of plots in which each genotype was observed. The denominator of equation 3 is equivalent to the phenotypic variance, 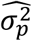. Standard errors of the estimated heritabilities were approximated using the delta method (Lynch and Walsh, 1998; Holland et al., 2003).

For each irrigation regime, genotypic 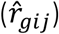 and phenotypic 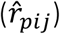 correlations between traits and their standard errors were estimated using multivariate REML in PROC MIXED of SAS version 9.4 (SAS Institute 2013) as previously described (Holland et al., 2001; Holland, 2006). To eliminate model convergence issues arising from the differences in scale among the various elements, a data standardization procedure was implemented using PROC STANDARD in SAS version 9.4. The BLUEs generated from Equation 2 for the two irrigation regimes within the individual years (e.g. 2010 WL, 2010 WW) were standardized to have a mean of zero and standard deviation of one prior to model fitting. The model used separately for each irrigation regime was as follows:

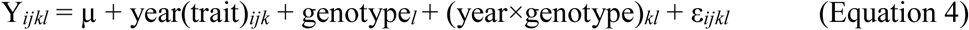

where Y_*ijkl*_ are the paired BLUEs for the *i*th and *j*th traits; μ is the multivariate grand mean; year(trait)_*ijk*_ is the effect of the *k* th year on the combined *i*th and *j*th traits; genotype*l* is the effect of the *l*th genotype; (year×genotype)*kl* is the effect of the interaction between the *k* th year and the *l*th genotype; and ε _*ijkl*_ is the random error term. The terms genotype*l* and (year×genotype)*kl* were fitted as random effects while year(trait)_*ijk*_ was fitted as a fixed effect. The REPEATED statement was used to estimate the covariance of the error associated with the *i*th and *j*th trait BLUEs measured for the same genotype.

The formula for estimating genotypic correlations was as follows:

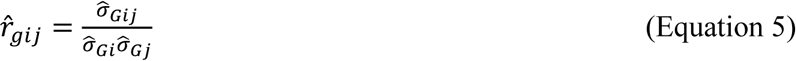

where 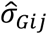 is the estimated genotypic covariance between traits *i* and *j*, 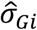 is the estimated genotypic standard deviation of trait *i* and 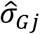 is the estimated genotypic standard deviation of trait *j*.

The formula for estimating phenotypic correlations was as follows:

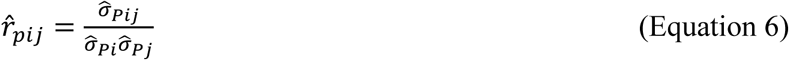

where 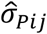 is the estimated phenotypic covariance between traits *i* and *j*, 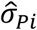 is the estimated phenotypic standard deviation of trait *i* and 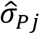 is the estimated phenotypic standard deviation of trait *j*. For both genotypic and phenotypic correlations, significance was assessed by computing the standard errors for the respective correlation values using the delta method based on Taylor series expansion (Lynch and Walsh, 1998; Holland et al., 2003). A correlation value greater than ±0.12 for either genotypic or phenotypic correlations corresponded to a confidence interval not including zero, thus designated as a statistically significant correlation (*P* < 0.05).

### QTL Analysis

The marker genotyping and genetic map construction for the TM-1×NM24016 RIL mapping population was previously reported in Gore et al. (2014). Briefly, 841 marker loci, consisting of 429 simple-sequence repeat (SSR) and 412 genotyping-by-sequencing (GBS)-based single nucleotide polymorphism (SNP) loci, were assigned to 117 linkage groups covering ~2,061 cM of the cotton genome. The 841 marker loci were not equally distributed across the genome. The placement of markers on the allotetraploid cotton (*G. hirsutum* L. acc. TM-1) draft genome assembly for marker-chromosome assignment is described in Pauli et al. (2016a).

#### QTL mapping with Bayesian classification method

We employed a Bayesian classification method to implement a multi-marker mapping technique as described in Zhang et al. (2005) and Zhang et al. (2008). Briefly, the goal of the analysis was to identify single markers that explained a significant amount of the variability for the seed elemental concentrations within each of the individual irrigation regimes. With a total of *n* observations (or RILs) and *m* markers for each observation (or RIL), let *i* = *1,2,…,n* be the index for each observation (or RIL) and *j =1,2,…, m* be the index for the markers across all linkage groups. Then, the phenotypic value for the *i*th observation (or RIL) within an irrigation regime, WL or WW, was modeled as follows:

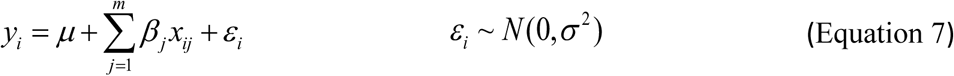

where *μ* is the overall mean, *x*_*ij*_ is the genotypic value of the *j*^th^ marker of individual *i*, and *ɛ*_*i*_is the random error term from the environmental factors. In this model, the parameter of interest is *β*_*j*_, representing the main effect of the *j*^th^ marker. Note that this model allows simultaneous identification of main effects using marker information from the entire genome.

Due to the large number of predictor variables in the model (i.e., markers) but relatively small number of observations available, the inference/estimation was done via a Bayesian framework using Markov chain Monte Carlo. The inference/estimation was implemented as a two-step procedure using a Gibbs sampler. *A priori* information was incorporated into the model by prior specification for the parameter of interest (spike and slab prior where marker effect is either positive, negative, or negligible), and estimation was based on the corresponding posterior distributions. In the first step, all main marker effects were ranked based on their posterior probabilities of having a non-zero effect on the trait of interest, and in the second step, a subset of all marker effects were selected. The number of effects selected in the first step depended on the total number of observations to ensure that efficient parameter estimates were obtained while keeping a desirable level of statistical power.

The BLUEs for each of the profiled elements were fitted independently within each of the irrigation regimes using Equation 7. In all analyses, the first 5,000 iterations were discarded as burn-in period and the following 5,000 iterations were used for inference/estimation. Model convergence was confirmed by the diagnostic tools presented in Cowles and Carlin (1996). For each parameter of interest, we estimated its magnitude and direction, as well as the posterior probabilities of being greater or less than zero. These posterior probabilities were used to calculate the Bayes factor as defined by Jeffreys (1935 and 1961). As advised by Jeffreys (1961), a Bayes factor between 10 and 100 provides “strong evidence” and larger than 100 means “decisive evidence.” Therefore, a QTL was declared significant if it had a Bayes factor ≥100.

#### Seemingly unrelated regression analysis

We implemented seemingly unrelated regression (SUR), a multi-trait (multivariate) analysis, to identify QTL controlling phenotypic variation in multiple elemental concentrations. As a generalization of linear regression models with multiple responses, SUR was first proposed by Zellner more than half a century ago (Zellner, 1962) and has been successfully applied to analyzing high-dimensional metabolite data sets (Chen et al., 2015; Chen et al., 2017). The method of SUR includes a set of multiple regression equations with each equation representing one of the response variables (i.e., a single elemental trait) of the multivariate response while assuming that the error terms are correlated between equations. For our analyses, elements were grouped according to their biological functions as described in Taiz and Zeiger (2006) and Mengel and Kirkby (2012), which created two groups with more than two elements. The “ionic” group (Ca, K, Mg, and Mn) included elements that remain in ionic form in plant tissue, and the “redox” group (Cu, Fe, Mo, Ni, and Zn) that contained elements involved in oxidation/reduction reactions. For each irrigation regime, WL or WW, the following model was fitted to each of the three groups of elements:

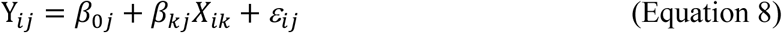

where Y_*ij*_ denotes the concentration of each of the individual *j*^th^ elements within the defined elemental groupings for the *i*^th^ RIL, *β*_*0j*_ represents the regression intercept, *β*_*kj*_ is the regression coefficient for the *X*_*ik*_ predictor that denotes the genotype of the *k* th marker of the *i*^th^ RIL, and *ɛ*_*ij*_ is the error term that follows 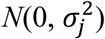 and cov(*ɛij ɛlk*). Such potential correlation structure among elemental traits allow the SUR model to have higher statistical power than linear regression models having only a single elemental trait. Note that for different RILs when *i* ≠ l, the term *σ*_*jk*_ is equal to zero. The data for each element were standardized to have mean zero and a standard deviation of one prior to the analysis, allowing regression coefficient estimates and their associated 95% confidence intervals for each element within a group to be comparable.

Nomenclature of QTL were defined by combining the elemental name, chromosome assignment, linkage group, and peak marker position of the identified QTL having names preceding with “q” to denote QTL. To declare QTL as mapping to the same location, we used a 10 cM window to be consistent with previous studies of this population (Pauli et al., 2016a).

### Prediction of Irrigation Regime

We used the elemental concentration BLUEs generated for individual years to predict whether genotypes were grown under WL or WW conditions in the three years this study was conducted. First, the BLUEs were standardized within each year to have a mean of zero and a standard deviation of one to account for the effect of each individual year (non-reproducible environments). Next, we employed a cross-validation strategy whereby two of the years were used to predict the remaining year, for example, BLUEs from 2010 and 2011 were used to predict irrigation regime in 2012. To assess the prediction accuracies, Pearson’s correlation coefficient was calculated between observed and predicted values. The various models tested in R (R Core Team, 2016) were as follows: logistic regression; linear discriminate analysis (LDA) and quadratic discriminate analysis (QDA) implemented in the *MASS* package (Venables and Ripley, 2002); k-nearest neighbors (KNN) implemented in the *class* package (Venables and Ripley, 2002); and support vector machines (SVM) implemented in the *e1071* package (Meyer et al., 2017).

To visualize and understand the relationship between irrigation regimes and the RILs, the BLUEs calculated separately for each year were used together to conduct a principal components analysis (PCA) using the “prcomp” function in the *stats* package (R Core Team, 2016). In contrast to the prediction of irrigation regime, the BLUEs were not centered and scaled prior to the PCA because this analysis was intended to explore the observed relationships among the yearly fluctuations in elemental concentrations.

### Data Availability

BLUPs calculated from the fitted mixed linear model for soil elemental concentrations are contained in File S1. BLUEs, both overall and by-year, for seed elemental concentrations are contained in File S2. Genotypic data for the 95 RILs from the TM-1×NM24016 mapping population with accompanying linkage map information are contained in File S3. File S4 contains the genetic linkage map information integrated with the published TM-1 draft genome sequence (Zhang et al. 2015).

## RESULTS

### Soil elemental variability

The soil samples taken from throughout the experimental field site demonstrated that spatial variability existed for the concentration of the 14 elements profiled (Supplemental Figure 1). A year effect of at least moderately high significance was only found for K, Rb, S, and Zn (*P* < 0.01, Supplemental Table 1), although the effect of depth at which soil samples were taken was highly significant for essentially all elements profiled. Because the year effect was weakly significant (*P* < 0.05) or non-significant (*P* > 0.05) for 10 of the 14 elements, both years of data were used in conducting geostatistical analyses to reveal and quantify the spatially structured and heterogeneous nature of the soil element concentrations across the field site (Figure 1). The fitted models produced effective ranges of spatial correlation from 12.49 to 66.44 m, with an average distance of 26.68 m (Supplemental Table 2). As expected, these results confirmed that concentrations of nearly all soil elements were related in a distance-dependent manner throughout the field. The only element that did not display any type of detectable spatial relationship was sulfur, which had an estimated concentration of 1089.90 ppm across the entire field site (Supplemental Table 3).

**Figure 1.**
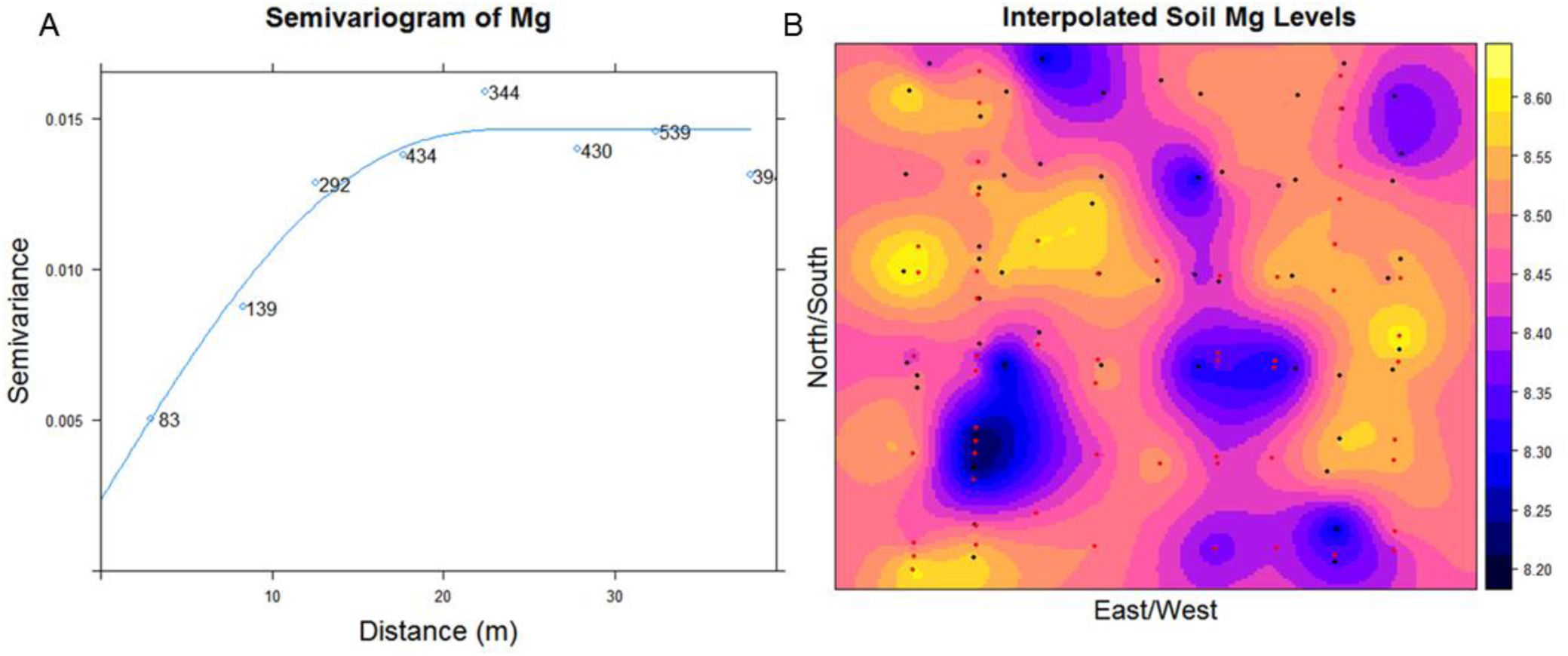
Characterization of soil magnesium (Mg) concentration in the field site where the mapping population was evaluated. A) Variogram representing spatial continuity of Mg variability; samples become spatially independent at a distance of 24.98 m. Values near the fitted line within the plot denote the number of point pairs at a given distance. B) Interpolated Mg concentrations (log transformed, parts per million) throughout the field. Black and red colored dots represent the sampling locations in 2010 and 2012, respectively.

The soil conditions were not limiting for plant growth as the concentration of key nutrients for cotton production in Arizona, which are Fe, K, Mn, P, and Zn all had minimum values exceeding production recommendations (Supplemental Table 3) (Silvertooth, 2001). Arsenic, an element that can be toxic for plants, had an observed maximum value of 4.99 ppm, which was well below the accepted threshold of 40 ppm (Walsh et al., 1977). Given the empirical evidence provided by the soil element analysis and the precision irrigation management used in this experiment, we are confident that high temperature and water deficit were the primary abiotic stresses impacting cotton plants from first flower to harvest.

### Seed ionome profiles

In order to understand how the seed ionome responds to abiotic stress, two irrigation regimes, consisting of water-limited (WL) and well-watered (WW) conditions, were imposed at flowering (50% of plots at first flower) and continued throughout the remainder of the season until harvest. Coinciding with the irrigation regimes, day time temperatures in the desert Southwest, on average, exceeded 32°C, the threshold above which lint yields are acutely impacted (Pauli et al., 2016a; Schlenker and Roberts, 2009). To control for the effects of localized soil environment and phenological development, the interpolated soil elemental concentrations and flowering time of genotypes were individually tested as covariates in the mixed linear model (Equation 2) to assess their association with seed element concentrations. The only elements that exhibited a significant (*P* < 0.05) linear relationship between seed and soil levels were Co, Mg, and Rb (Supplemental Table 4). The seed element concentrations that had an association (*P* < 0.05) with flowering time were As, Cu, and Ni (Supplemental Table 4). After accounting for the effects of soil environment and phenology, we observed significant genotypic differences (*P* < 0.0001) for all 14 elements; however, only seven of the elements, Ca, Cu, Fe, Mg, Mo, S, and Zn, displayed differences between the irrigation regimes. Of these seven elements, only the concentration of Mg decreased under WL conditions, while the concentrations of the other six elements increased under WL conditions. Genotype-by-irrigation regime interactions were only significant for Co, Mn, Mo, and S (Figure 2, Supplemental Table 4).

**Figure 2.**
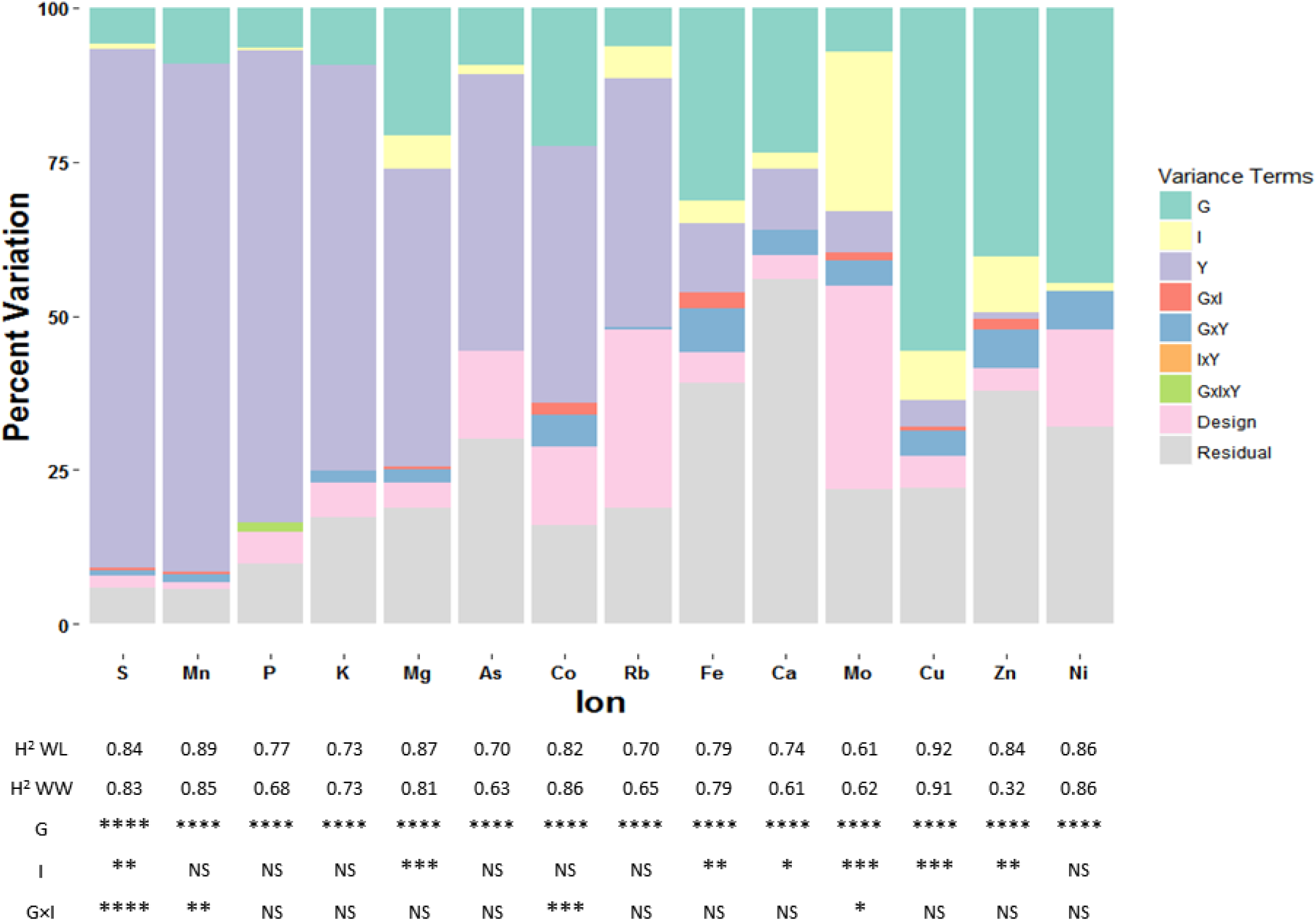
Sources of variation for cotton seed elements. The figure shows the decomposition of phenotypic variance into respective components: teal for genotypic (G), yellow for irrigation regime (I), purple for year (Y), red for genotype-by-irrigation regime interaction (G×I), blue for genotype-by-year interaction (G×Y), orange for irrigation regime-by-year interaction (I×Y), green for the three way interaction of genotype-by-irrigation regime-by-year (G×I×Y), pink for field design variables replication, block, and column, and gray for residual variance. Variance component estimates were calculated from modeling all terms in Equation 2 as random. The table below lists the broad-sense heritabilities (*Ĥ*^*2*^) for the two irrigation regimes, water-limited (WL) and well-watered (WW), and the significance of the *P*-values for the different fixed effects from Equation 2. **** *P*-value < 0.0001, *** *P*-value < 0.001, ** *P*-value < 0.01, * *P*-value < 0.05, and NS > 0.05.

In terms of the relative seed elemental concentration values, which were weight normalized and rescaled prior to analysis, the RIL population exhibited extensive phenotypic variation for the 14 elements profiled (Table 1). The macronutrients Ca, K, Mg, P and S all had average concentrations above 10,000 ppb with K being the element in highest concentration with values over 46,000 ppb in both irrigation regimes. Arsenic was the element with the lowest relative average concentration of only 10.58 and 10.53 ppb for WL and WW conditions, respectively, and Co was the second least abundant element with averages of 12.09 and 12.12 ppb for WL and WW regimes, respectively. Both positive and negative transgressive segregation were observed for most elements in the RIL population.

**Table 1.** Summary statistics of cotton seed elements. Means, standard deviations, and ranges (parts per billion) of best linear unbiased estimators (BLUEs) for seed elements for the TM-1×NM24016 recombinant inbred line (RIL) population evaluated under two irrigation regimes, water-limited (WL) and well-watered (WW) conditions, including parental lines and their midparent values. Field trials were conducted from 2010-12 at the Maricopa Agricultural Center located in Maricopa, AZ.

| Ion | Irrigation regime | Parents |  |  | RIL population |  |  |  |
| --- | --- | --- | --- | --- | --- | --- | --- | --- |
|  |  | TM-1 | NM24016 | Midparent | Mean | Std. Dev. | Min. | Max. |
| As | WL | 10.60 | 10.54 | 10.57 | 10.58 | 0.09 | 10.43 | 10.88 |
|  | WW | 10.57 | 10.54 | 10.56 | 10.53 | 0.07 | 10.39 | 10.74 |
| Ca | WL | 12528.86 | 17067.21 | 14798.04 | 13125.79 | 2357.68 | 8316.97 | 18824.00 |
|  | WW | 11163.35 | 14774.05 | 12968.70 | 12206.20 | 2125.10 | 6348.93 | 16899.59 |
| Co | WL | 12.29 | 13.09 | 12.69 | 12.09 | 0.38 | 11.40 | 13.03 |
|  | WW | 12.29 | 12.92 | 12.60 | 12.12 | 0.42 | 11.12 | 13.08 |
| Cu | WL | 33.42 | 94.22 | 63.82 | 61.58 | 10.65 | 33.27 | 93.37 |
|  | WW | 30.74 | 86.60 | 58.67 | 55.56 | 10.41 | 26.29 | 80.54 |
| Fe | WL | 237.12 | 459.06 | 348.09 | 305.44 | 50.25 | 210.46 | 427.10 |
|  | WW | 212.06 | 412.20 | 312.13 | 279.95 | 46.66 | 175.77 | 388.16 |
| K | WL | 46164.35 | 48992.02 | 47578.18 | 46143.43 | 3280.07 | 38486.65 | 54048.57 |
|  | WW | 47709.49 | 49919.47 | 48814.48 | 46905.04 | 3529.59 | 40055.63 | 55012.74 |
| Mg | WL | 14729.61 | 18466.01 | 16597.81 | 16797.41 | 1934.97 | 11709.99 | 20884.97 |
|  | WW | 16230.02 | 20191.12 | 18210.57 | 18003.52 | 1863.47 | 13340.41 | 21434.30 |
| Mn | WL | 112.29 | 162.72 | 137.50 | 121.36 | 14.22 | 87.42 | 152.30 |
|  | WW | 105.08 | 148.24 | 126.66 | 118.26 | 14.49 | 81.94 | 161.27 |
| Mo | WL | 19.02 | 21.00 | 20.01 | 19.74 | 1.24 | 17.32 | 22.76 |
|  | WW | 16.86 | 19.44 | 18.15 | 17.52 | 1.00 | 15.23 | 19.85 |
| Ni | WL | 13.21 | 14.28 | 13.75 | 14.11 | 0.72 | 12.50 | 15.52 |
|  | WW | 12.88 | 14.00 | 13.44 | 13.88 | 0.77 | 12.25 | 15.92 |
| P | WL | 45776.14 | 52804.11 | 49290.12 | 51054.12 | 4184.02 | 41247.43 | 61589.54 |
|  | WW | 45192.32 | 55452.35 | 50322.34 | 52830.91 | 3838.64 | 40742.95 | 63115.53 |
| Rb | WL | 28.77 | 29.79 | 29.28 | 30.02 | 2.17 | 24.87 | 34.98 |
|  | WW | 26.75 | 27.05 | 26.90 | 27.67 | 1.95 | 22.97 | 33.41 |
| S | WL | 14526.02 | 15488.89 | 15007.46 | 14663.85 | 1306.94 | 11394.59 | 18026.07 |
|  | WW | 14087.15 | 15004.32 | 14545.73 | 13937.97 | 1276.18 | 10665.15 | 16948.30 |
| Zn | WL | 208.40 | 357.09 | 282.74 | 275.27 | 44.09 | 160.72 | 357.82 |
|  | WW | 171.51 | 303.45 | 237.48 | 248.19 | 42.92 | 139.85 | 328.92 |

We estimated broad-sense heritabilities for the elements to determine the extent to which phenotypic variation was attributable to genetic variation in the RIL population. Heritability values were moderate to high, ranging from a minimum of 0.32 (Zn, WW conditions, Figure 2) to a maximum of 0.92 (Cu, WL conditions). With the exception of the low estimate for Zn under WW conditions, heritabilities were all greater than 0.60. With regard to the heritability estimates between irrigation regimes, there were no significant differences (two-sided *t*-test, *P* > 0.05). The ANOVA also revealed that the variance due to year effects was large for most elements (> 40% for As, Co, K, Mg, Mn, P, Rb, and S) and that the variances associated with the second-and third-order interaction terms were small (Figure 2).

To characterize the relationship among the elements and potentially shared regulation of seed elemental levels, we estimated pairwise phenotypic 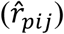 and genotypic 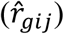 correlations among the 14 elements. Element pairs with strong phenotypic correlations under both irrigation regimes (ranging from 0.51 to 0.77) included Mg/P, Mg/Zn, Mg/Fe, Ca/Mn, Fe/Zn, and Zn/P, results in agreement with other plant studies (Baxter et al., 2013; Zhang et al., 2014; Shakoor et al., 2016). Neither major differences in terms of element pairings nor contrasts in correlation strengths were observed between the two irrigation regimes (Figure 3, Supplemental Table 5). Phenotypic correlations were, on average, positive with the exception of As, which was negatively correlated with all other elements (Figure 3). The most highly correlated elements, Ca, Cu, Fe, Ni, Mg, Mn, P, and Zn, were grouped together at the center of the network, while micronutrients As, Co, Mo, and Rb were mostly on the perimeter (Figure 3). Interestingly, K was one of the least correlated elements despite being a macronutrient. It was primarily correlated with Rb, the only other monovalent cation profiled, suggesting that chemical similarities among the elements are responsible for their relatedness. The genotypic correlations estimated for the TM-1×NM24016 RIL population closely followed the pattern observed for the phenotypic correlations with respect to strength and pairings (Figure 3, Supplemental Table 5 and 6).

**Figure 3.**
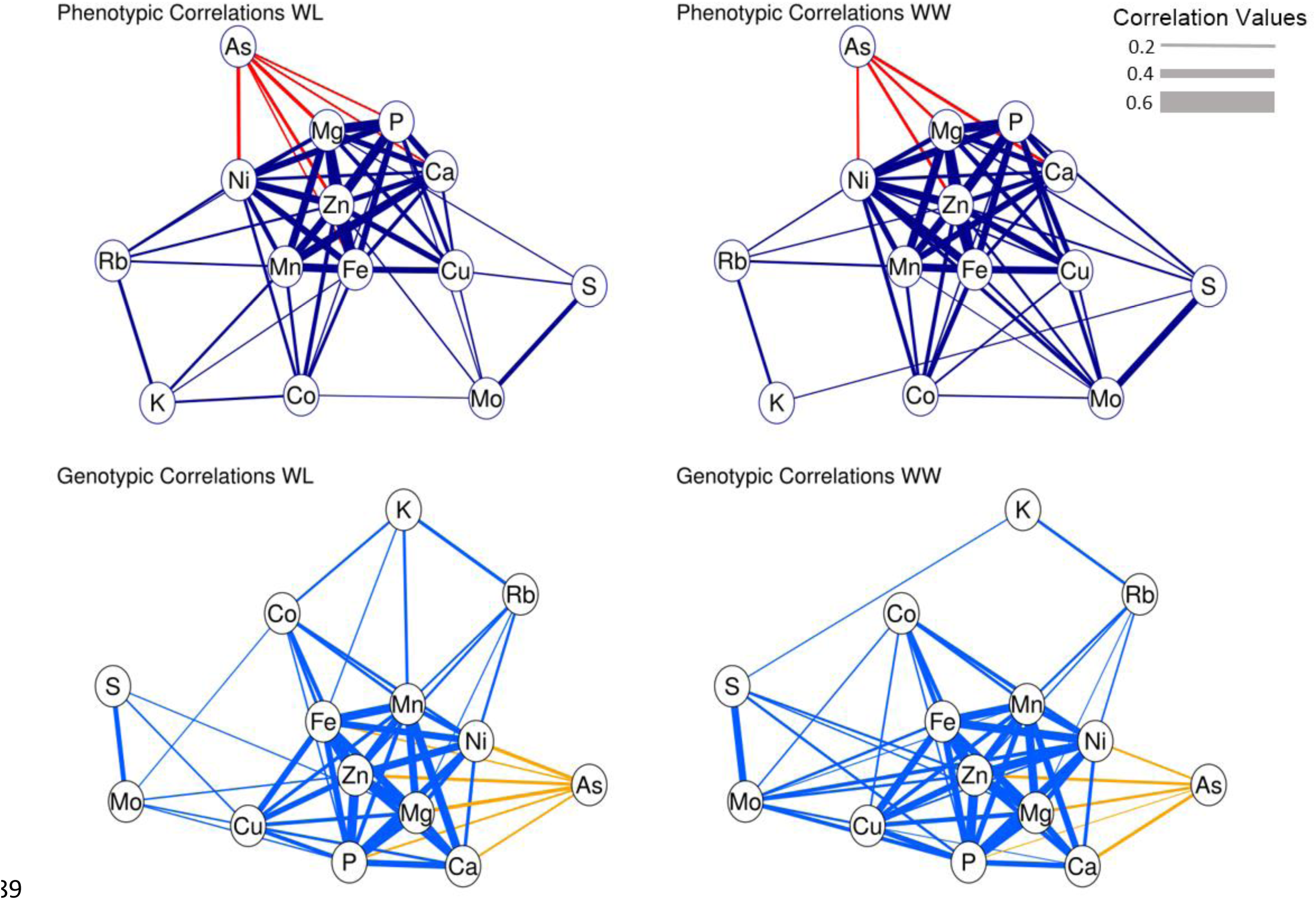
Network correlation graph of phenotypic 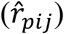 and genotypic 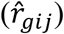 correlations among elements profiled in cotton seed. Purple and blue edge colors represent positive correlation values, red and gold represent negative correlation values. The edge thickness represents the magnitude of the correlation value with only those values greater than 0.20 being displayed (*P* < 0.05).

### QTL mapping

The mapping of QTL utilizing a Bayesian classification method detected a total of 38 QTL that mapped to 15 chromosomes and 21 unique genomic locations (Figure 4, Supplemental Table 7). Concerning the two irrigation regimes, 16 and 22 QTL were detected under WL and WW conditions, respectively. Only four QTL, *qCu.A07.28.00*, *qMg.A12.45.21*, *qNi.D12.116.00*, and *qP.A05.74.05*, were detected for both irrigation regimes. The number of QTL found per element varied from one (Mo) to five (Mg and Rb). There were a total of five genomic regions, located on chromosomes A05, A06, A12, D01, and D12, to which two or more QTL mapped with all but A12 having ion pairs (Ca/Mg, Fe/Zn, Cu/Ni, and Fe/Ni, respectively) with similar biochemical properties. Several QTL mapped to chromosome A05, linkage group 74, including for Fe, Mg, Ni, P, and Zn, indicating that this genomic region has a significant impact on the cotton seed ionome.

**Figure 4.**
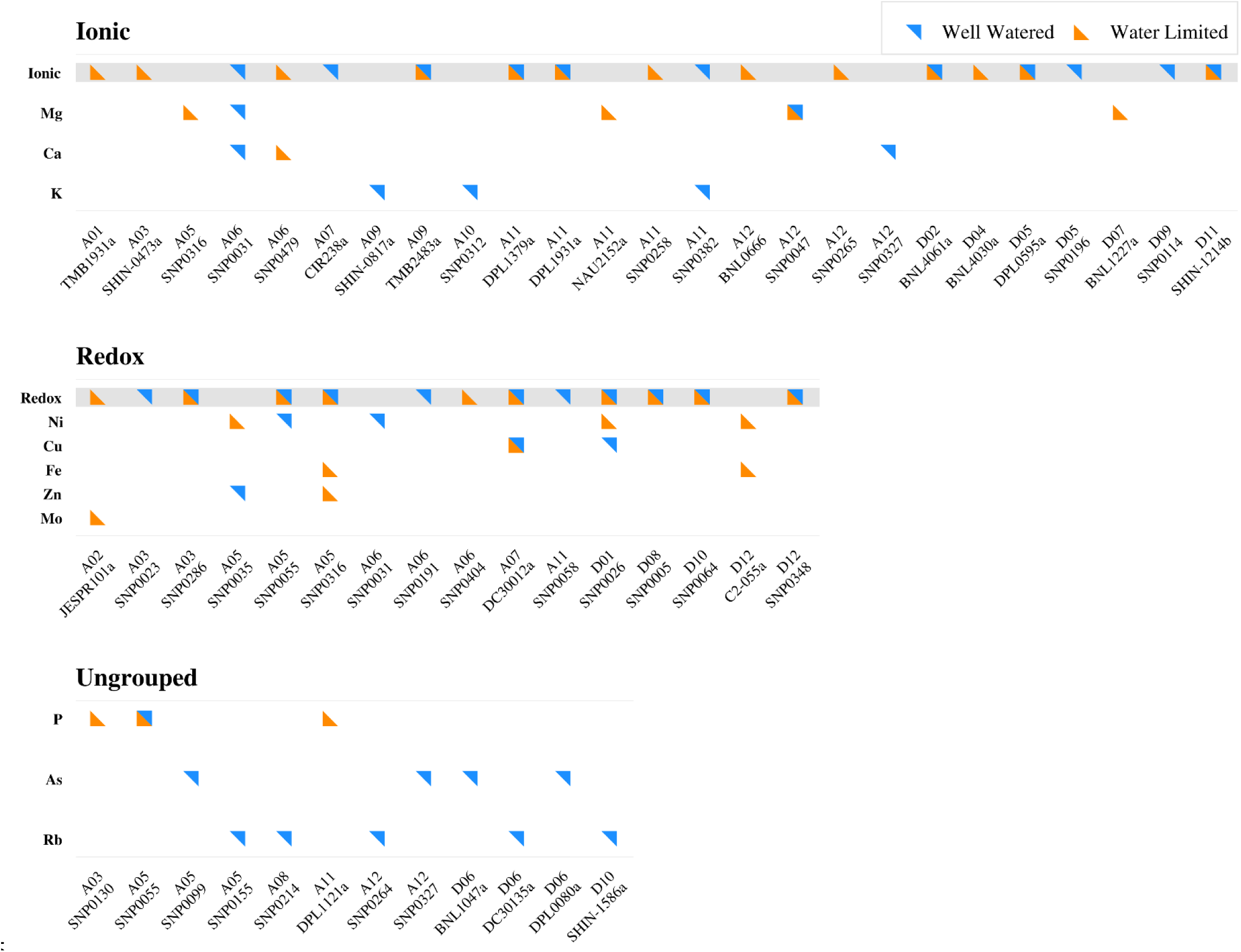
Identified QTL controlling phenotypic variation for elemental concentration in cotton seed. Results from the multi-trait analysis, in which elements were grouped into ionic and redox categories, are highlighted in gray. The results for the individual elements analyzed using the Bayesian classification are listed below the corresponding multi-trait groupings. Loci that were shared across ionic and redox groups are highlighted with symbols above the respective chromosome and marker names. “Ungrouped” denotes individual elements that were not contained in the multi-trait groupings and analyzed using only the Bayesian classification method. No QTL were detected for Co, Mn, and S, thus these elements are not shown.

We also implemented a more statistically powerful multi-trait mapping approach (a multivariate analysis) to reveal QTL controlling phenotypic variation for elements with similar properties. Elements were first grouped based on their biochemical function creating two groups of elements: “ionic” and “redox.” Both of these groups were then analyzed using the method of seemingly unrelated regression (SUR) to identify QTL impacting multiple elemental concentrations in cotton seed. This multi-trait analysis identified 45 QTL that mapped to 18 chromosomes and 31 unique genomic locations (Figure 4, Supplemental Table 8). Nearly an equal number of QTL were detected for each irrigation regime, with 23 and 22 QTL found for the WL and WW irrigation regimes, respectively. In contrast to the Bayesian mapping approach, which only detected four QTL for both irrigation regimes, the multi-trait analysis detected 14. Additionally, the multi-trait analysis also identified 10 QTL that, although detected with the Bayesian classification approach, were not declared significant because they were below the Bayes factor threshold of 100 (Supplemental Table 9).

We examined the tetraploid cotton draft genome sequence (*G. hirsutum* L. acc. TM-1, which was one of the parents of the RIL population) (Zhang et al., 2015), to determine if there were plausible candidate genes underlying detected QTL. For QTL that mapped to A06, A07, and D12, candidate genes found to colocalize included a copper transporter, metal tolerance protein, and potassium transporter, respectively. For the Zn QTL on A05, which was detected by both mapping methods as well as for both irrigation regimes, a zinc transporter gene described in *G. hirsutum* (UniProt ID K9N1X9) was identified within the 342 kb interval defined by the flanking markers. The Bayesian classification-detected QTL for potassium on A09, *qK.A09.33.00*, defined a 1.9 Mb interval containing a gene involved in metal ion transport (UniProt ID B9IFR1).

### Prediction of irrigation regime

We conducted a principal component analysis (PCA) on best linear unbiased estimators (BLUEs) for all 14 elements, revealing that the first two PCs accounted for almost half (47.3%) of the total elemental variance. The PCA revealed a distinct separation between the two irrigation regimes, with PC 2 on the y-axis largely separating the WL and WW into two groups and explaining 14.1% of the total variance (Figure 5A). These results suggested that the seed ionome could be predictive of abiotic stress. To test this hypothesis, five supervised machine learning approaches, logistic regression, linear discriminate analysis (LDA), quadratic discriminate analysis (QDA), k-nearest neighbors (KNN), and support vector machines (SVM), were used to determine the irrigation regime within which a RIL was grown. To help control for non-reproducible environmental effects, the elemental BLUEs for the individual years were first centered and scaled within respective years prior to their use in the five models. Extremely high prediction accuracies were obtained for all five methods, but SVM achieved the highest average prediction accuracy across the three years at 97.7% (Figure 5B) and a maximum accuracy of 98.5% for 2011. The KNN method produced the lowest average prediction accuracy of 92.5%, which was observed in 2010 and 2011. The remaining three methods, LDA, logistic regression, and QDA, had an across-year-and-irrigation regime average of ~94%.

**Figure 5.**
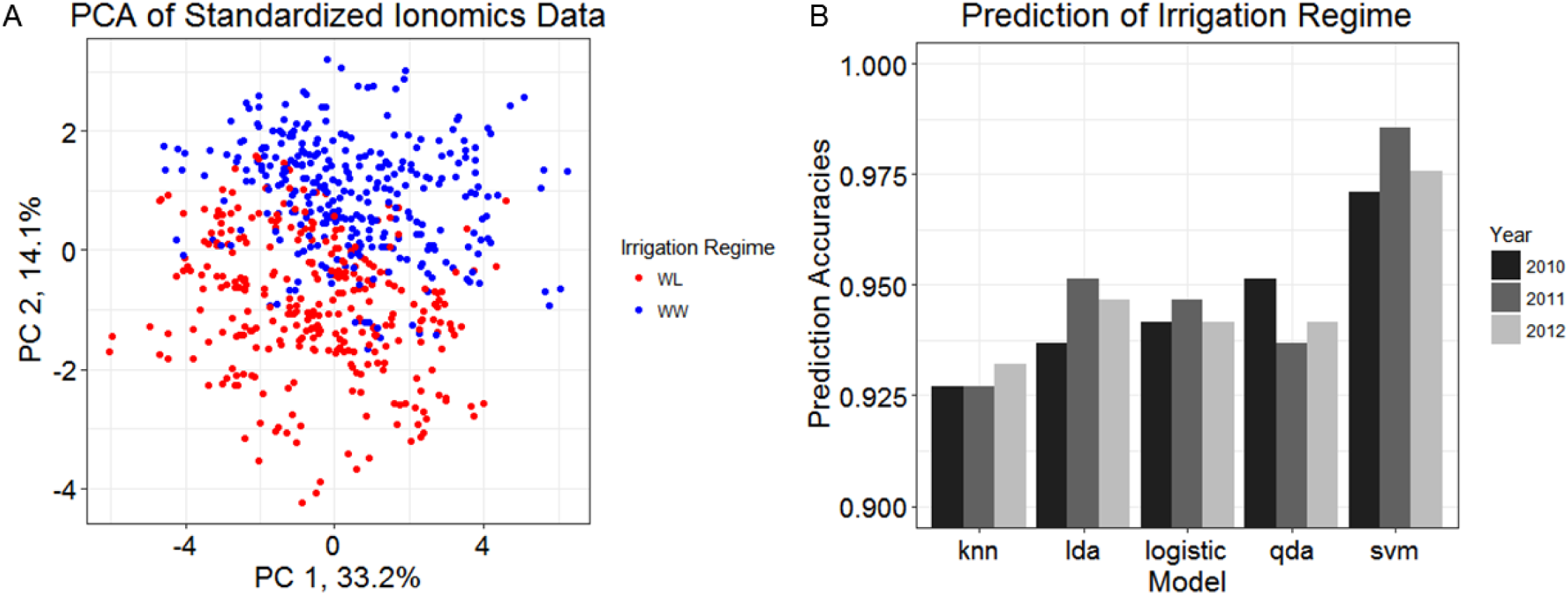
Characterization and prediction of irrigation regime using seed ionomic data for the 14 elements. A) Results of a PCA using standardized individual year BLUEs from 2010-12 for the 95 RILs. Colors indicate the irrigation regime in which RILs were evaluated. B) Prediction accuracies for irrigation regime achieved using five different classification methods. Bars represent across-year cross validation accuracies.

## DISCUSSION

Over the last decade, ionomics has been established as a powerful tool for both examining the nutrient status of plants to assess homeostasis and for revealing the genetic mechanisms responsible for elemental variation. However, research efforts have largely been focused on characterizing the elemental concentration of various plant tissues and identifying mutant lines for further genetic characterization (Lahner et al., 2003; Baxter et al., 2009; Pinson et al., 2015). These studies have led to valuable knowledge on the genetic control of element accumulation in plants, but have offered limited insight into how the ionome interacts with the environment. To address these information gaps in ionomics research, we evaluated a cotton RIL mapping population under contrasting irrigation regimes to assess the effects of water deficit on the ionome in a hot, arid environment. The elemental profiles of the localized soil environment were also analyzed so that these results could be incorporated into the analysis of the cotton seed ionome. Further expanding on this work, the elemental concentration data were utilized for prediction of abiotic stress, as defined by WL or WW irrigation regimes, to investigate how accurately the ionome predicts the physiological status of the plant.

The heterogeneous nature of the soil environment can impact phenotypic variation of most quantitative traits and influence growth characteristics like root density, biomass allocation, and interplant competition (Caldwell et al., 1996; Hutchings et al., 2003). Because of this innate relationship between the soil environment and plant development, we used geospatial interpolation methods to model the elemental concentrations across the field site (Figure 1) in order to assess if there was a direct association between soil and seed element levels. Although we only found three elements (Co, Mg, and Rb) that exhibited a significant linear relationship between soil and seed concentrations (Supplemental Table 4), under other more varied environments the associations may be stronger and more numerous among the elements. Without the inclusion of the soil data it would have been more difficult to determine if variation observed in the seed ionome was due to genetic effects, abiotic stress, or only the localized environment. This was demonstrated in the case of Mg, which had both a significant irrigation effect and a linear relationship between soil and seed Mg concentrations. The inclusion of the interpolated soil level data permitted us to decouple the impacts of water deficit stress from soil variability and improve our genetic mapping, along with calculation of phenotypic and genotypic correlations.

The irrigation regimes imposed in this experiment provided the ability to evaluate how the cotton seed ionome responds to abiotic stress, specifically water deficit. Seven of the 14 elements assayed had significant differences between the irrigation regimes (Figure 2), including Ca, Mg, and S which are important macronutrients (Mengel and Kirkby, 2012). Although we can only speculate what mechanisms may be responsible for observed differences in most element concentrations due to water deficit, the increased seed Ca concentration under WL conditions is consistent with its involvement in intra-plant signaling and osmoregulation via increased solute concentration. This hypothesis is in agreement with the results of Patakas et al. (2002) who found similar elevated Ca levels in leaves of grape (*Vitis vinifera* L, cv. Savatiano), another woody perennial species like cotton, when evaluated under drought-stress conditions. Although their analyses were based on leaf tissue samples and not seed, one could hypothesize that seeds would show a similar response based on Ca signaling, which occurs both as an early and secondary signaling response, with local and global effects mediated through transport in the xylem (Knight, 1999; Sanders et al., 1999; Xiong et al., 2002; White and Broadley, 2003). Additionally, the elevated levels of Ca in seeds harvested from WL plots could also be due to increased levels of Ca associated with stomatal closure in response to drought stress (Atkinson, 1991; Schroeder et al., 2001; Taiz and Zeiger, 2006). However the long-term dynamics of Ca flux in these responses and what influence they would have on the seed ionome remains unclear given that time-course studies are currently lacking in the literature.

To gain insight into the ionome and potential joint regulation of elemental accumulation in cotton seed, we evaluated the correlations, both phenotypic and genotypic, among the elements profiled (Figure 3). The phenotypic correlations confirmed our initial hypothesis that elements with similar biological relevance and chemical properties would be highly correlated, consistent with the results of Shakoor et al. (2016) and others. However, the lack of a contrast in the correlation values and patterns between the two irrigation regimes was somewhat surprising (Figure 3). Initially, we hypothesized that differences in available soil moisture would impact the relationship among the individual elements. To evaluate if this association was due to environment or genetics, we assessed the genetic correlation among the elements using a multivariate restricted maximum likelihood approach, an analysis not previously used in ionomic studies. The results of these analyses mirrored those obtained for the phenotypic correlations, namely that the strength and relationship among genotypic correlations were highly similar across the two irrigation regimes. In both sets of correlations, the majority of macronutrients clustered in the center of the correlation network graphs, along with those micronutrient elements involved in redox reactions (Cu, Fe, Ni, and Zn) (Taiz and Zeiger, 2006; Mengel and Kirkby, 2012). The remaining elements were grouped along the perimeter including arsenic, which was the only element to be negatively correlated with all other elements, most likely due to its toxicity to plants and thus exclusion (Meharg and Hartley-Whitaker, 2002). Although our study represents an ideal situation given the precisely controlled water deficit stress and single field site, the consistent trends in correlation, both phenotypic and genotypic, support the supposition that the cotton ionome is a highly interrelated system under strong genetic control.

Given the observed genotypic correlation values suggesting a shared genetic basis responsible for elemental accumulation, QTL mapping was carried out to assess if loci responsible for phenotypic variation were indeed shared amongst the elements. A Bayesian classification method (Zhang et al., 2005) that fitted all markers simultaneously while exploiting *a priori* information was used to more fully control for the genetic background effects to enable better detection of causal loci. This approach was successful in detecting QTL, but more importantly it identified six genomic regions to which multiple QTL mapped. The QTL for elements that colocalized to the same location, such as Ca/Mg on A06, Fe/Zn on A05, and Ni/Fe on D12, had similar biochemical properties, such as being divalent cations, and were also involved in parallel biochemical processes like regulation of osmotic potentials and electron transfer (Taiz and Zeiger, 2006).

To date, most ionomic genetic mapping studies, whether linkage analysis or genome-wide association studies, have relied on univariate mapping approaches (Baxter et al., 2013; Baxter et al., 2014; Zhang et al., 2014; Asaro et al., 2016; Shakoor et al., 2016). Despite the ability of these methods to detect QTL that individually contribute to phenotypic variation, they fail to account for the relationship among the various elements, and thus the shared biology underpinning these traits (Baxter, 2015). Given the QTL results from the Bayesian analysis in which QTL impacting physiologically related elements colocalized to genomic regions, and the shared genetic basis revealed by genotypic correlations, a novel analysis was needed to more fully capitalize on the inter-trait relatedness.

With these considerations in mind, a multi-trait (multivariate) mapping approach was taken to exploit the relationships among the elements to improve the ability to detect QTL controlling the accumulation of multiple elements within the cotton seed. Seeming unrelated regression (SUR, Zellner 1962) was implemented so that elements with similar characteristics could be grouped together and treated as one phenotype. The colocalizing QTL from the Bayesian analysis corroborated the division of elements into “ionic” and “redox” groups posited in the literature (Mengel and Kirkby, 2012) and further supported the use of a multi-trait analysis. By analyzing like elements in aggregate and accounting for the correlation among them, statistical power to detect QTL was increased. This improved power led to more consistent detection of QTL with respect to irrigation regime; 14 QTL were identified in both WL and WW conditions compared to four QTL found in the Bayesian analysis. These results are more congruent with what the genetic correlations revealed, largely that relationships among elements are stable despite the perturbations by abiotic stress. Additionally, the multi-trait analysis detected 10 QTL whose Bayes factors were below the significance threshold in the Bayesian analysis, and thus missed, further highlighting why previous genetic mapping studies not utilizing a multi-trait analysis likely missed identifying important loci. Although the two mapping approaches rely on two distinctly different branches of statistics, Bayesian and frequentist methods, there was agreement between the two; a total of 18 QTL were concordant between methods providing further support for the mutually identified QTL (Figure 4). Also, elements with co-located QTL had similar ionic charges, suggesting that the genetic factors underlying these QTL are not element-specific but instead dependent on chemical properties like those used by cellular ion transporters for ion selectivity (Tester, 1990).

Although the detected QTL explained a moderate amount of the phenotypic variation observed for cotton seed elemental concentrations, there was a question of whether these data themselves could describe the physiological status of the plant. The results from PCA clearly demonstrated that the elemental data could capture the effects of water deficit and served to separate the RILs into respective irrigation regimes in which they were evaluated (Figure 5A). Building on these results, various supervised machine learning algorithms were used to predict which irrigation regime RILs were evaluated under out of the three years in which this experiment was conducted. When the year-to-year variation was removed via standardization, extremely high prediction accuracies, greater than 92%, were achieved for all of the methods used with minimal variation in prediction amongst years within a method. These results demonstrate that the ionome is capable of encapsulating the physiological status of cotton plants without the use of more traditional physiological phenotypes like carbon isotope discrimination, relative leaf water content, osmotic adjustment, and other, more time-consuming and costly measurements.

## CONCLUSION

The ionome is capable of capturing the mineral and nutrient content of the plant tissue from which samples are taken thereby offering valuable insight on the physiological status of the plant (Salt et al., 2008). Because of this, ionomic profiling was used for both cotton seeds and the ambient soil to study how the ionome interacts with and responds to its localized environment, with a focus on the impact of water deficit. Our results provide further evidence that the ionome is a complex, interrelated biosystem that is largely under shared genetic control, and as such, it responds as an integrated unit to abiotic stress as evident by the stable relationship among phenotypic and genotypic correlations despite the contrasting irrigation treatments. Because of the interrelatedness among the ionomic traits, a genetic mapping approach that capitalizes on this shared genetic architecture was used to detect loci that influence the composition of the various elements in plant seed. To further extend these findings and concepts, the ionome was found to have a remarkably high predictive accuracy for irrigation regime status that accurately reflected the physiological status of the plant. Although this was a two-category classification problem of drought-stressed versus non-drought stressed, it reflects the ability of these types of data to describe the status and overall wellbeing of the plant. Taken together, the present work provides new insight into how complex biological systems are controlled at the genetic level through multiple shared loci that are associated with correlated responses, including responses to water deficit regimes.

## ACKNOWLEDGEMENTS

Mention of trade names or commercial products in this publication is solely for the purpose of providing specific information and does not imply recommendation or endorsement by the USDA. The USDA is an equal opportunity provider and employer. We especially thank Kristen Cox, Bill Luckett, Joel Gilley, Virginia Moreno, Sara Wyckoff, Spencer Fosnot, Brian Nadon, Kimberly Green, and Melissa Jurkowski for providing excellent technical expertise. We would also like to thank Kelly Thorp and members of his lab for managing the soil samples. A special thanks to Dan Ilut for graphical assistance and review of the manuscript. Thanks to Christine Diepenbrock for critical review of the manuscript.

